# Tropical high-altitude insects show limited capacity to handle high temperatures

**DOI:** 10.1101/2022.09.10.507406

**Authors:** Harshad Vijay Mayekar, Pooran Singh Solanki, Homica Arya, Rajaguru Aradhya, Prashanth Suravajhala, Volker Loeschcke, Subhash Rajpurohit

## Abstract

Growing summer season and increased anthropogenic activities pose a continual challenge to resident species. Ectotherms like insects are especially vulnerable to rapid climatic changes. High-altitude tropical insect populations have been rarely examined for their responses to high-temperature. We exposed a tropical out-bred highland population of *Drosophila melanogaster* from the Himalayas to growing summer conditions in outdoor mesocosm units. Population response to thermal changes was tracked over ninety days at both phenotypic and genotypic level. Whole genomic resequencing data suggested a clear seasonal shift in allele frequencies. Interestingly, the general heat responsive genes were missing in the summer due to monsoon allele shift; an atypical response noted for high-altitude tropical populations. Instead, candidates involved in kinases and phosphorylation emerged as key players. Heat-knockdown time decreased over time indicating a limited ability to handle increasing temperature. Merging data from both allelic shifts and heat-knockdown time indicated a limited capacity for high-altitude insects in coping with climate warming.

## 1. Introduction

Temperature is a dominant climatic variable with overarching effect on organismal fitness (Stevens 1989). Rising global temperatures raise alarms of extinction for several species which are unable to adapt to climate change at shorter time scales (Turney, Ausseil et al. 2020, Habibullah, Din et al. 2022). Rising temperatures aside, variability in temperatures is also a confounding factor increasing the complexity of assessing the impact of climate change (Duffy, Gouhier et al. 2022) as diverse zones face different climatic regimes. Temperature rise in tropics was considered to be lower than in temperate or polar regions, recent climate models strongly predict tropical zones of Amazonia, Sahel, South-east Asia and India to face increased warming (Holmes, Woollings et al. 2016, Bathiany, Dakos et al. 2018). In Amazonia the effect could be predominantly due to soil drying, but in the Asian tropics e.g. India, it is largely attributed to variation in atmospheric forces. Not surprisingly, the Indian monsoon patterns (Menon, Levermann et al. 2013), have varied greatly in recent years; with recurring floods supporting the correlation with rising temperatures. Yet, conclusions from climatic models could be taken with a caveats since tropical mappings are scarce (Bathiany, Dakos et al. 2018). Even then the global impact of climate change on living organisms cannot be ignored (Root, Price et al. 2003). In India, the increased incidence of heat-waves (1300 people died in 2010 in Ahmedabad alone (Knowlton, Kulkarni et al. 2014)) has been predicted to rise in several parts of north, central and western India with annual temperatures expected to rise from 2.2^0^C to 5.5^0^C (Kumar, Wiltshire et al. 2013, Dholakia, Mishra et al. 2015). Rising temperatures could be detrimental to survival of organisms and need research attention.

Anticipating thermal breadths could perhaps provide cues to organismal responses to rising temperatures. Janzen (1967) surmised broader thermal tolerance for temperate species than tropical species since the tropics have evolutionarily experienced stable thermal ranges. Hence seasonal effects are generally less pronounced in the tropics. However this does not include places of high elevation/ altitude from the tropics which have cooler temperatures compared to lowlands. Tropical altitudinal zones are distinct with little spatial overlap. Hence, populations from higher altitudes could be expected to have responses different from lowlands. Whether high altitudinal tropical populations mimic temperate/ polar zone patterns or display a different pattern is not known as tropical species have been scarcely studied for their response to climate change (Feeley, Stroud et al. 2017).

Ectotherms regulate their body temperatures based on the external environment (Hertz, Huey et al. 1993) and are ideal to study thermal responses to changing climate and several species are known to be impacted by climate (Duffy, Gouhier et al. 2022). Insects as notable ectotherms; are seen to exhibit tremendous populations declines (Halsch, Shapiro et al. 2021). Coping with climate change has been tested across several insects (Sears et al., 2011; Woods, 2013), but none parallel the diverse trends observed in the vinegar fly, *Drosophila melanogaster*. *D. melanogaster* likely originated in Sub-Saharan Africa (Lachaise, Cariou et al. 1988) and shortly attained a global distribution (David and Capy 1988). Within a few ten thousand years, the sub-tropical *D. melanogaster* populations adapted worldwide to spatially diverse climatic selection pressures, e.g. alcohol tolerance (Berry and Kreitman 1993, Fry, Donlon et al. 2008), diapause (Schmidt, Zhu et al. 2008), circadian rhythm (Svetec, Zhao et al. 2015), thermal stress and tolerance (Weeks, McKechnie et al. 2002), (Fallis, Fanara et al. 2014), and metabolic load (Sezgin, Duvernell et al. 2004). However, whole genome sequencing offers more insights into climatic adaptations in *D. melanogaster* rather than interpretations based on few genes, loci or makers (Turner, Levine et al. 2008, Fabian, Kapun et al. 2012, Adrion, Hahn et al. 2015). Surprisingly, genomic analyses revealed parallel trends in adaptation in north American and Australian *D. melanogaster* populations compared to their southern counterparts respectively (Reinhardt, Kolaczkowski et al. 2014). Not only spatial but also temporal heterogeneity due to changing seasons could affect adaptation to climate change (Bergland, Behrman et al. 2014, Rudman, Greenblum et al. 2022). Thermal variation could be experienced not only at the annual scale, but also at the diel and within and across generations or even at the seasonal scale (Wang and Dillon 2014, Kefford, Ghalambor et al. 2022). Typically, tracking thermal performance curves (Sinclair, Marshall et al. 2016) or investigating traits exhibiting plastic responses to thermal variation (Seebacher, White et al. 2015, Sgrò, Terblanche et al. 2016) have been instrumental in understanding climate change. However, temporal genomic studies indicate seasonally changing allele frequencies are predictable and adaptive (Bergland, Behrman et al. 2014, Machado, Bergland et al. 2021). It should however be noted that most of the spatial and/ or temporal genomic studies of *Drosophila* thermal adaptations stem from temperate zones (Rudman, Greenblum et al. 2022).

Tropical organisms living closer to their upper thermal limits could find the pace of temperature rise challenging (Sørensen, Kristensen et al. 2016). Temperate *D. melanogaster* populations respond to cold hardening through phenotypic plasticity (Stone, Erickson et al. 2020). Whether tropical ectotherms respond to thermal fluctuations through phenotypic plasticity or local adaptation is not clear (Ghalambor, Huey et al. 2006). Phenotypic plasticity could either buffer responses (Muñoz 2022) to climate change or even accelerate adaptive responses (Harshman and Hoffmann 2000). Yet whether, plastic changes keep pace with variable climatic patterns needs attention(Kapun and Flatt 2019). Limited capacity to handle thermal stress has been hinted irrespective of latitude or elevation (Sunday, Bates et al. 2014). Yet, a broad geographic sampling of insect populations from the tropics and their response patterns to climate change are currently not known, especially from the Asian continent (particularly South-east Asian region). Besides, high altitudinal tropical insect populations have rarely been addressed for genomic changes in response to climate change.

Here, we investigate the genomic and physiological response of a high altitudinal tropical *D. melanogaster* population from Western Himalayas to heat stress across a growing summer season. *D. melanogaster* populations collected were outbred under laboratory conditions, released into the mesocosm units in a tropical orchard set-up at the Ahmedabad University Experimental Ecology and Evolution station with overlapping generations, and sub-sampled at three different time points (TP’s) (spanning a growing summer season with a duration of ninety days. Samples were subjected to whole genome re-sequencing and we wanted to look for genomic responses to warming i.e. changes in thermal responsive genes (TRGs) (Venkatachalam and Montell 2007, Matsuura, Sokabe et al. 2009). Since *Drosophila* piggybacks between these thermal preferences, particularly in the early life stages, state of relative conservation among species and unrelated temperature changes at species origin was felt necessary to understand the TRGs oscillations if any. At the physiological level, we tested the heat knock-down times of populations collected from the three time points across the summer season. We aimed to investigate the macro physiological pattern of a cold-adapted high altitudinal population to a tropical growing summer.

## 2. Materials and Methods

### 2.1 *Drosophila* collection and maintenance

*D. melanogaster* population was collected from higher altitudinal orchards from Western Himalayas (Rohru, Himachal Pradesh, India; 31.2046^0^N, 77.7524^0^E, Altitude 1554 meters) using banana baits in the year 2018. Each collected female was placed in a separate tube (iso-female line) at the site of collection. Almost 270 such lines were made in the field using females captured during the field trip. Upon 10 to15 hour’s arrival to the lab, vials were placed at room temperature (25^0^C). Successfully established lines were examined based on the taxonomic keys to reconfirm their identity at the species level. Overall, 40 lines of *D. melanogaster* were successful and were maintained in the laboratory, on standard ‘agar-jaggery-yeast-maize’ media at 25^0^C, with a 12/ 12light/dark cycle. All the 40 lines were used in this work.

### 2.2 Out-crossing

Forty isofemale lines of *D. melanogaster* were outcrossed (founding numbers 800 adults: 400 males + 400 females) in a 0.61m X 0.61m X 0.61m dimension cage made up of plexiglass acrylic sheet. For this, 10 males and 10 females each from 40 isofemale lines were used. Each generation of eggs was collected in twenty half-pint size culture bottles. As soon as the pupae became darker and ready to eclose, bottles were moved to the plexiglass cage and plugs were removed. This allowed random mating in the founding base population. Once all the flies had enclosed in the cage, eggs were collected and transferred to fresh food bottles for the next generation. These lines were outcrossed for 5 generations keeping populations’ size between 5-6 thousand individuals every generation.

### 2.3 Mesocosm Cages

To track adaptive changes in real time, and whether or not changes in temperature (increasing temperature over growing tropical summer) lead to changes in allelic frequencies, we tracked populations (3-fold replication) in outdoor mesocosm cages over a period of 90 days (Fig. 1). Outdoor (mesocosm) cages were set up at the Ahmedabad University Experimental Evolution Study Station. Each mesocosm was a 1.5x1.5x1.5 meters dimension (custom designed at Ahmedabad University Fabrication Shop) outdoor insect rearing enclosure with a mature (dwarf) Sapota tree. Three cages were used for this experiment (MESO-1, MESO-2, and MESO-3). Each cage was founded with 1000 males and 1000 females collected from the 6^th^ generation of laboratory cage (i.e. outcrossed population). Every morning at 09.00 am fresh food (50 ml of standard ‘agar-jaggery-yeast-maize’) in a half-pint bottle was placed in each enclosure for the entire duration of the experiment (15^th^ April 2020 to 15^th^ July 2020). Flies were allowed to oviposit on the fresh food for 24 hours. Each morning, bottles with eggs laid in the last 24 hours, were sealed with cotton plugs and larvae were allowed to develop (inside the same cages); upon eclosion, bottles were opened and adults were released into the cages. Thus populations were cultured under a natural regime of overlapping generations. Temperature and RH in the cages were recorded using HOBO U23 Pro v2 data loggers (Onset Computer Corp., Bourne, MA, USA).

**Figure 1.**
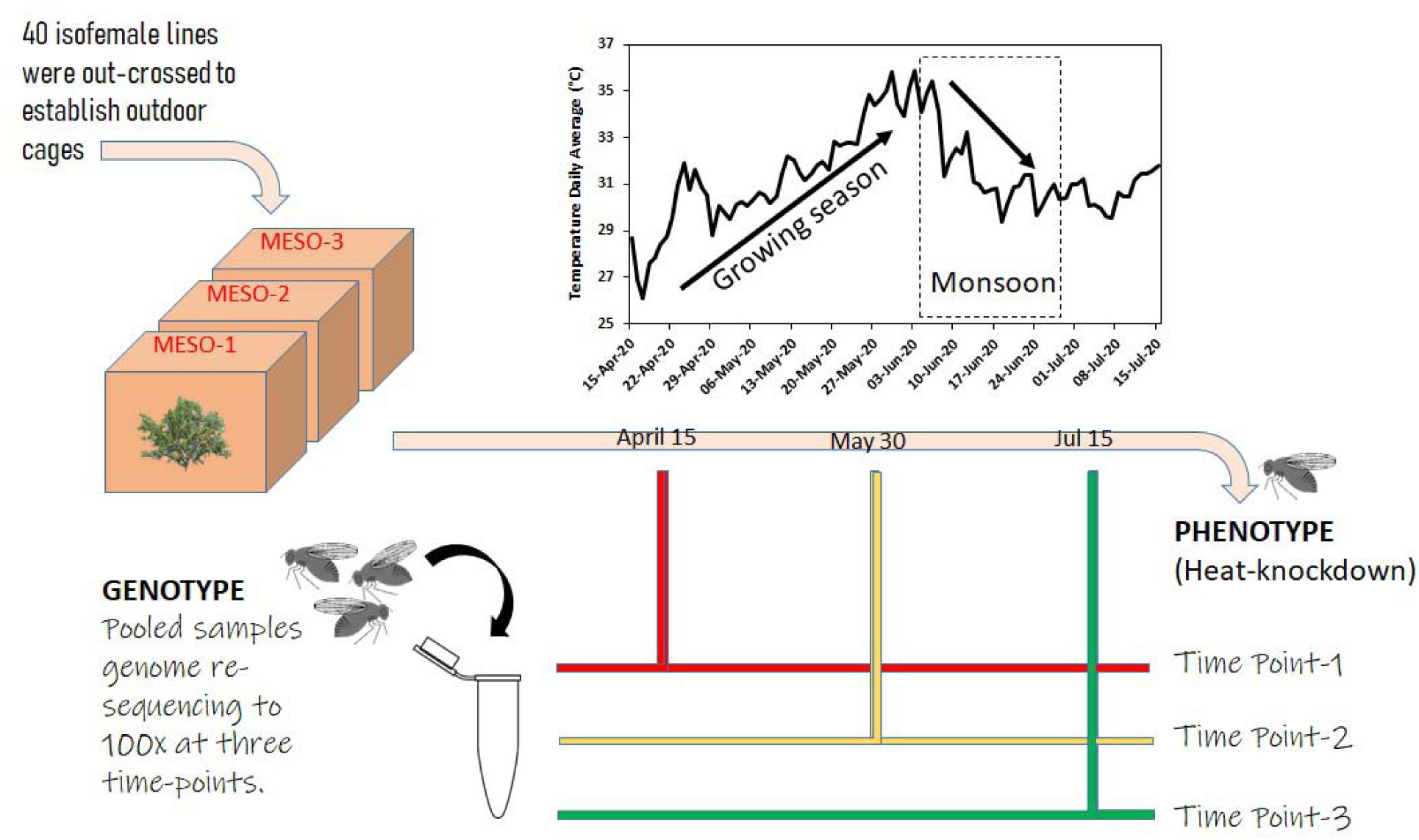
Outdoor mescocosm experimental set-up. Forty *D. melanogaster* isofemale lines were pooled to generate an outbred population. Lines were outcrossed for 5 generations in 24x24x24 inch dimension cages under laboratory conditions (25^0^C). From this outcrossed population (maintaining the size close to 5000-6000 individuals every generation) we released 500 males and 500 females in each outdoor mesocosm cages (MESO-1, MESO-2, and MESO-3). For the next 90 days food was changed every day and the previous bottle with eggs was capped and placed in the cage itself. Upon eclosion adults were released in the existing cage and bottles were discarded. Cages were sampled at three time-points (TP) for genomic and phenotypic analysis: at the beginning (TP1), in the middle (TP2) and at the end (TP3).

### 2.4 Adaptive tracking of populations

Samples for genomic and phenotypic measurements were collected at three time-points for adaptive tracking. Samples for TP1 were collected from the founding adults while the TP2 and TP3 samples were collected on the 45th and 90th day, respectively. Eggs were collected in 30 bottles at lower density (10 bottles for each replicate cage). These bottles were allowed to develop under a constant temperature (25°C) in the laboratory and phenotypic data was collected using F3 individuals.

### 2.5 DNA sample preparation for genomic analysis

Pooled genome re-sequencing was performed with 100 *D. melanogaster* females in triplicate from each outdoor cage, a total of nine samples were collected for whole genome sequencing analysis. The DNA extraction procedures involved homogenizing the flies (n = 100) in 200 µL of lysis buffer (100 mM Tris-Cl, 100 mM EDTA, 100 mM NaCL, 0.5% SDS). The homogenate was incubated at 65°C for 30 minutes. Following this, proteins were precipitated by adding 800 µL of 2 parts 5M potassium acetate, 5 parts of 6M lithium chloride solution and 15 minutes ice incubation. The supernatant was collected after centrifugation of the homogenate at 12K RPM for 15 minutes at room temperature. DNA was precipitated by adding 800 µL of isopropanol and centrifugation at 12K RPM for 15 minutes. The pellet was collected and washed with 70% ethanol. After removing the ethanol, the pellet was allowed to dry at room temperature and was re-suspended in 100 µL of TE buffer.

### 2.6 Library construction and quality control

1μg of DNA was used as input material for the DNA sample preparations. Sequencing libraries were generated using NEBNext® DNA Library Prep Kit following manufacturer’s recommendations and indices were added to each sample. The genomic DNA was randomly fragmented to a size of 350bp by shearing, then DNA fragments were end polished, A-tailed, and ligated with the NEBNext adapter for Illumina sequencing, and further PCR enriched by P5 and indexed P7 oligos. The PCR products were purified (AMPure XP system) and resulting libraries were analyzed for size distribution by Agilent 2100 Bioanalyzer and quantified using real-time PCR.

### 2.7 Next Generation Sequencing

The whole genome sequencing (WGS) of natural populations was done to check the role of mutations in functional candidates. Library preparation and tagmentation was done using DNA-350 bp by default, while the sequencing was performed on a HiSeq PE150. The samples were mixed for library construction using a PCR-free library preparation guide. The downstream analysis and annotation was done with raw reads run through an in-house benchmarked pipeline (Meena, Mathur et al. 2018), tweaked for a whole exome pipeline (https://github.com/prashbio/WES).

After the allele frequencies were mapped, we read all mutation positions in MESO-1, MESO-2 and MESO-3 for all the three time point scales (T1|T2|T3) and created a data structure which contained three types of positions: single, double and triple mutations. For example, single types of mutations are those that are positioned only once across the 3 different TPs, double-any two TP scales while triple – across all three of them. This was done for MESO-1, MESO-2 and MESO-3 (experimental scale) and the resulting files were checked for interpreting the possible SNPs. However, we asked whether mutations that are identical are present at least at 2 time points at the same position and finally a composite matrix containing mutational positions of MESO-1, MESO-2, and MESO-3 was done.

### 2.8. Heat-knockdown assay

We tested F3 females (ca. 3 days old) for their thermal tolerance to an increasing temperature using the dynamic heat ramp assay. In this assay, temperatures ecologically relevant to the species are the base-point after which they are steadily increased till the fly is knocked down (Lutterschmidt and Hutchison 1997). Individual flies were collected in glass tubes of 0.05m^3^ volume. The tubes were immersed in a water bath set at room temperature (ca. 25^°^C).

Temperature was increased at the rate of 1°C/ minute with the upper thermal limit being 43^°^C. We measured three technical replicates with 24 flies each from three cages across three time points. Nested ANOVA (experimental cages within time-points) was used to assess the variation in knock-down time. Post-hoc analysis was performed using the *glht* () function from the “multcomp” package (Hothorn, Bretz et al. 2008). For genomic analysis, the flies were aspirated from the cages itself.

## 3. Results

### 3.1. Allele frequencies

The MESO-1, 2, & 3 replicates were run through the whole genome sequencing (WGS) pipeline where we inferred causal SNPs across these sub-populations (Table 1). The minor allele frequencies were tabulated from the depth with number of reads showing variation (DP4) with 1) forward ref alleles; 2) reverse ref; 3) forward alt; 4) reverse alt alleles, used in variant calling using the formula, variant allele frequency(VAF) = (forward alt + reverse alt alleles) / (forward ref alleles + reverse ref + forward alt+ reverse alt alleles). After setting up the MAF<=0.15 assuming that 15% of the SNPs would have this MAF, we further retained heterozygous SNPs. All 9 raw reads yielded 9,963,857 SNPs per sample. The most common SNPs between the MESO-1, 2, & 3 cages and the outliers were tabulated by mapping the chromosomal positions to flybase reference genome (flybase.org last accessed date: September 22, 2021) (Table 1/Fig. 2). By removing those specific minor allele frequencies (MAFs) above 0.15, we could assume that there is a clear distribution of contamination. We therefore sought to assess the possibility of observed TRG with MAF <0.15 which gave us an average of 240 SNPs per sample (Supplementary File 1). This raises the possibility that allele frequency could be generated by *D. melanogaster* alleles consistently at various time point scales. We further assessed the frequency of SNPs that are fixed for alternate alleles and cross checked this with the previously identified SNPs associated with seasonal changes (Temperature) and tabulated all SNPs for time points (Table 1).

**Figure 2.**
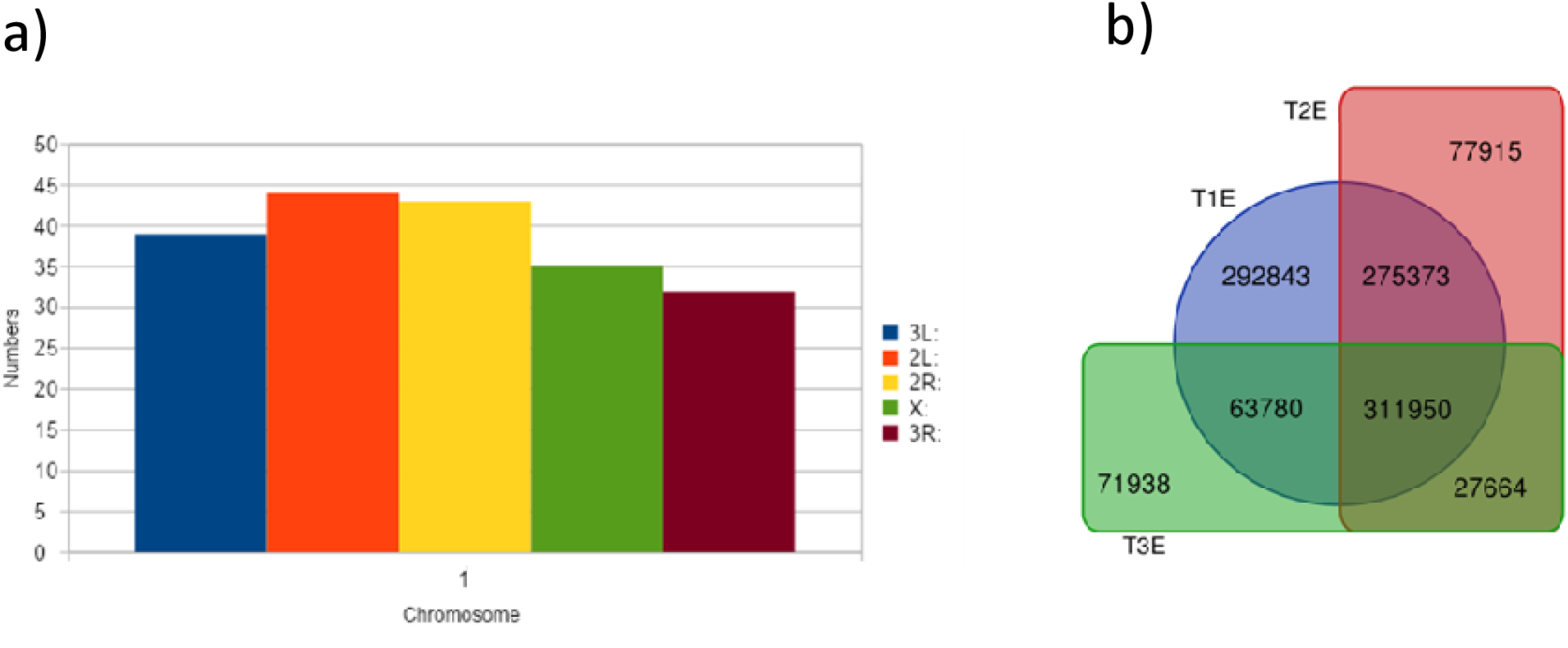
(a) Flybase SNPs mapped to our experimental candidates across all chromosomes (b) The SNPs mapped across the three time point experimental pools. The numbers indicate the individual time point SNPs apart from the most common SNPs at the intersection.

**Table 1:** Average number of SNPs after filtering.

|  |  |
| --- | --- |
| Total identified SNPs/ | 89,674,715/9,963,857 |
| Average SNPs per TP sample |  |
| $\geq 10$ DP $\leq 400$ | 5,015,244 |
| Total number of SNPs with MAF $<0.15$/ | 3,573,280/397,031 |
| Average MAF $<0.15$/sample | |
| Total used in analysis/ | 5,015,244/ 240 |
| Average bona fide SNPs per TRG |  |

### 3.2. Positional mapping with Flybase SNPs

The downstream annotation was processed with chromosomal alias obtaining 1870 total number of scaffolds with an average length of 17,195,935 per chromosome. The chromosomal distribution of SNPS is largely at the interface of 2L and X (Data not shown; Supplementary File 2 1, Fig. 3). We also mapped the SNPs to that of reported 193 SNPs from flybase to check if TRG mutations exist. We found Single TP SNPs: (n=noPOS: 4,666), Double TP SNPs: POS: (n=2,536) and no Triple TP SNPs (n=POS: 0).The final list of protein coding genes were mapped across various chromosomes (Supplementary File 1). As the current analysis represented identifying common mutations, we further looked into mapping the existing known SNPs (for heat stress).

**Figure 3.**
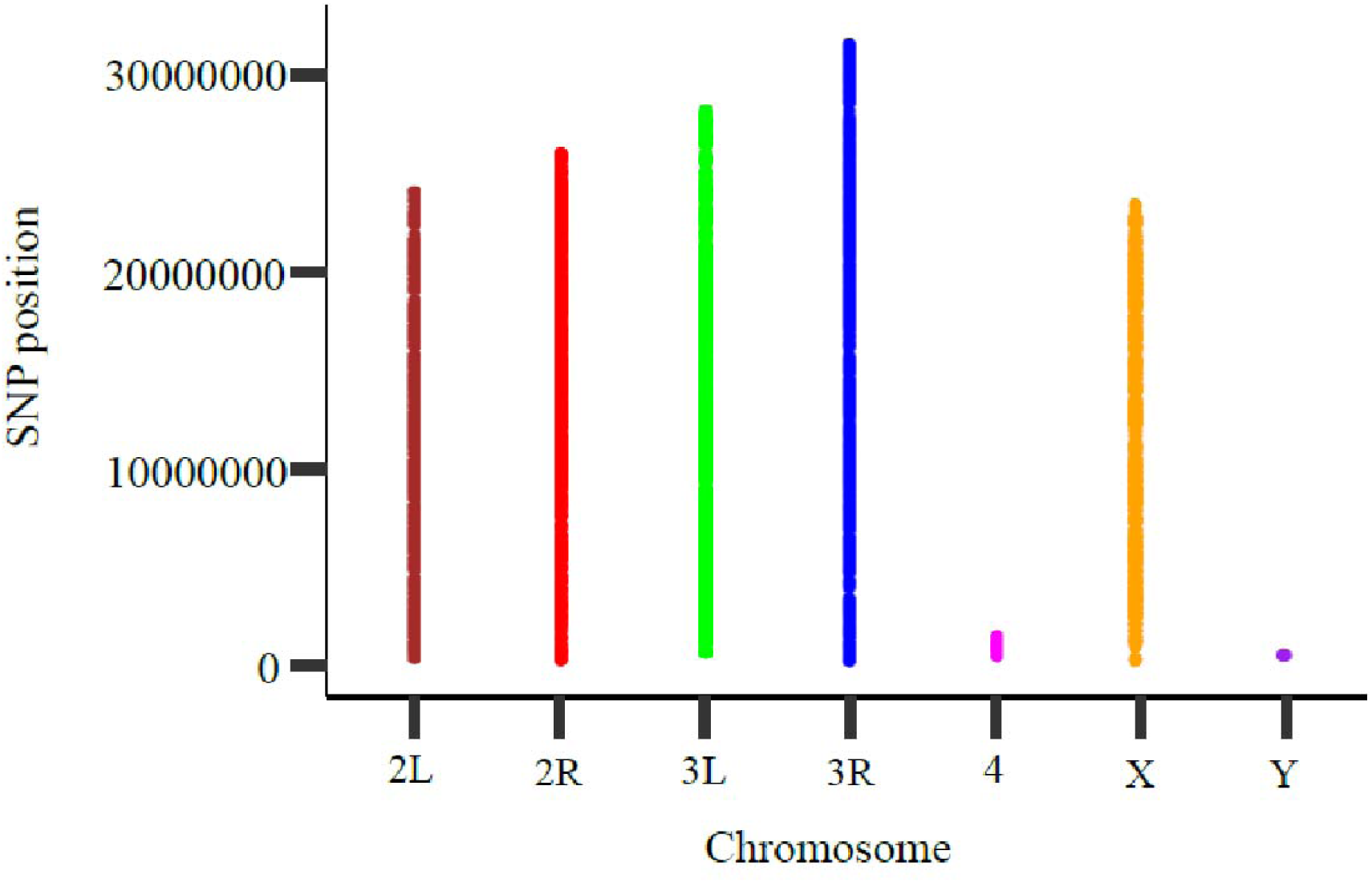
Variable polymorphisms shown to have a large deviation from time point scale 1 to time point 3 taking into account read depth and the number of sampled chromosomes. The X-axis shows the chromosomes with the Y-axis showing the single nucleotide polymorphism (SNP) positions.

### 3.3. Population comparisons

The time point samples and replicates mapped to the temperature variation across season regime were treated as T1_MESO-1, T2_MESO-1 and T3_MESO-1 as populations. As we obtain the data, the coverage across these three sub-populations were filtered for common mutations or SNPs that have been retained from T1_MESO-1,2&3 through T3_MESO-1,2&3 or the unique set of SNPs through them (Supplementary File 1). While removing the SNPs with total counts with the same allelic proportion, we deem that they have not undergone any changes through the three experiments. On the contrary, those SNPs that have an overlap with changes in SNPs across the same positions were filtered and we therefore limited the downstream analysis to such overlapping SNPs from three sub-populations or experiments. As we combined all the reads, we have checked the significant differences between the experiments for overlapping SNPs using the chi-square test as a standard practice. The *p-*value heuristics resulting from the analysis based on three population comparisons were in agreement. Another approach was to check the SNP mean density which could be done using an existing method (proposed by Burke et al (2010) and Winbush and Singh (2021) which measures the quantile score but our study is not a measure of nucleotide diversity from a threshold value instead the number of SNPs emerging from a whole genome sequencing study.

While we have found SNPs that have been deleted from T1 to T3 (Supplementary File 3), the MAFs in earlier cases were set to as big as MAF<0.89 which is 89%. Our stringent cutoff (Winbush and Singh 2021) indicates that highly sensitive SNPs emerge from our analysis and when these were tried to map to flySNP base, none or not many of them are mapped indicating that these are novel (data not shown). Nevertheless, we lacked sequencing data on similar scales where whole genome sequencing (WGS) was done so that we could have identified convergent or divergent regions contributing between the three sub-populations. For each comparison, we observed that 77,915 (24.97%), 71,938 (23.06%) and 27,664 (8.8%) SNPs in the respective three time points existed indicating that the temperature affects the sensitivity of the flies. We then mapped these SNPs to summarize the genes that have escaped the mutations in T3_MESO-1, 2 &3. When we map them to the table depicting counts for genes associated with these SNPs for T3_MESO-1, 2 & 3, we observe that a large number of diseased phenotypes are associated with these conditions. However, despite consistent read depth of coverage and SNPs, we observe a partial overlap suggesting that the accorded convergence can be attributed to slightly reduced heterozygosity which may invite a bias. However, this was reconfirmed from our validation set of founder experiments.

As we identified the genes with diseased phenotypes we were interested to see whether or not the candidate loci of these genes have an influence on the pathways. To check this, our panther gene ontology annotation derived enriched genes unique to temperature. In summary, our analysis identified a large number of candidate genes and SNPs associated with heat handling wiring machinery. Rather, our approaches indicate the variation across different subpopulations, as we argue that several diseased phenotypes have emerged from our analysis. Taken together, temperature sensitivity plays a very important role and serves as a determinant to mesocosm experimental study. Although genetic drift augurs well for taking temperature as a context, functional validation for observational variation is presumptive for maintaining such selection pressure. It may be argued that the function of the genes stemming from allelic variation has allowed us to determine candidate TRGs.

To check genomic signatures affected by seasonal fluctuations, we pooled sequencing data containing the SNPs by estimating the allele frequencies (AF). From each population and time point, the three replicate cages were merged and the files were segregated to average read depth of 20–200× coverage (Supplementary File 1). We observed that these estimated AFs have been shown to be accurate for understanding the magnitude of genetic variation through the time point scales. From Fig. 2, we reason that variable polymorphisms had a large deviation from time point scale 1 to time point 3 taking into account read depth and the number of sampled chromosomes. Of the 3,119,540 common SNPs tested, we identified substantial numbers falling between the statistical values, meaning they are called integrated thermal responsive SNPs. However, the AFs corresponding to selection coefficients per generation were not done with this statistical power as they are variable. Our rationale was to assess evolutionary features underlying rapid adaptive states in response to selection pressure. As the data is variable with the random disturbance and is different across elements of the vector, we asked whether the SNPs are enriched among functional genetic elements. To find out whether or not the SNPs are genic, we screened total SNPs present on specific genes common to these time points. A putative interaction map indicates that the genes are largely spread across with the key pathways attributing to these factors are MAP kinases and other important signaling pathways (Fig. 4). In general, we find four important genes and a few un-characterized genes which could be attributed to pleiotropy and linkage disequilibrium largely due to TRGs/seasonal variation.

**Figure 4.**
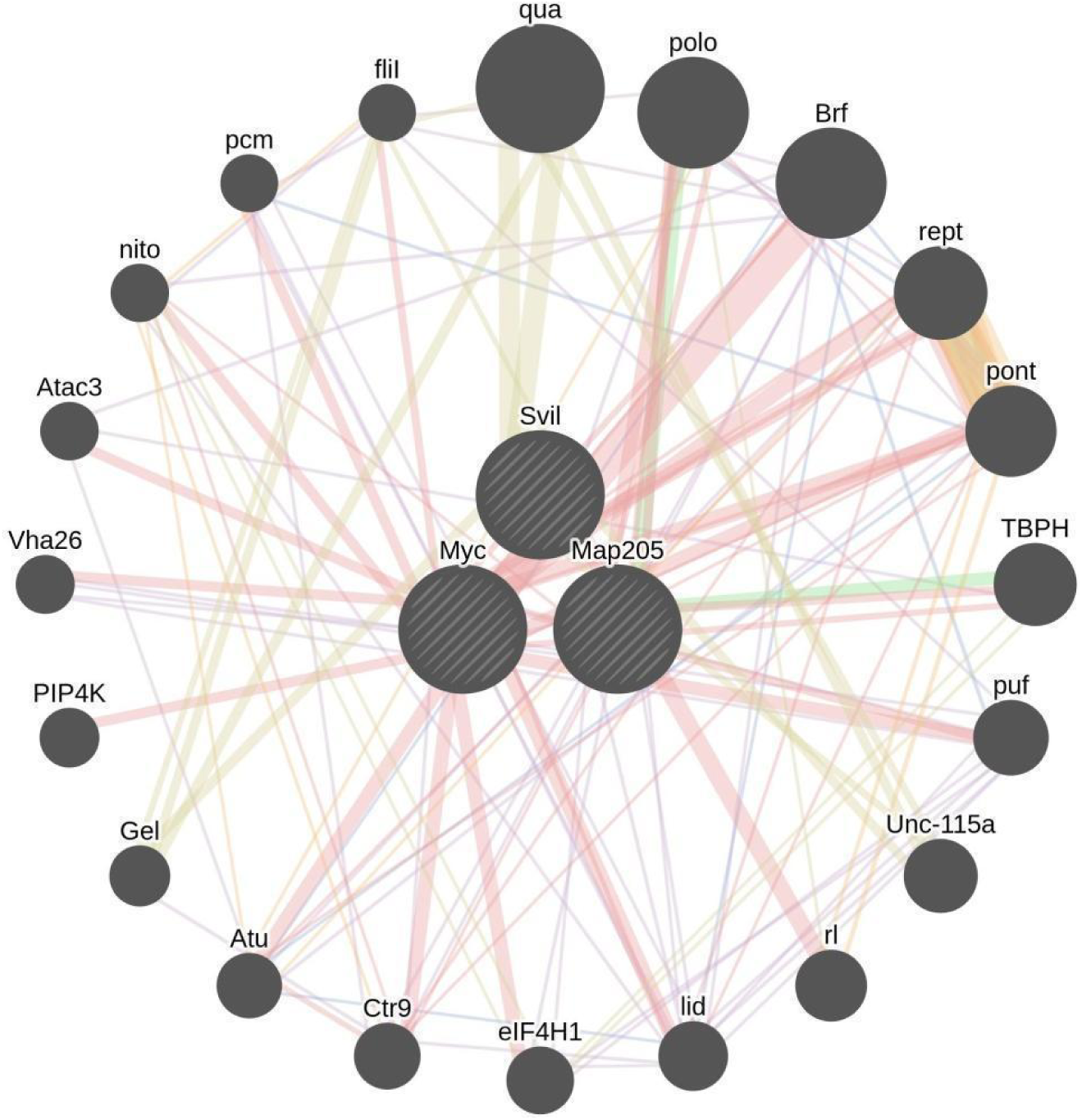
A putative interaction map of *Svil*, *Myc* and *MAP205* with interacting partners connected through edges (lines). The pink edges indicate that they are known to be physically interacting with each other while other edge colors denote pathways, domain association, co-localization, neighborhood and genetic interactions if any.

### 3.4. Heat tolerance

Knock-down times of flies significantly differed among all three time-points (df = 2, F= 966.30, P < 0.0001) (Supplementary File 4). Heat knockdown time was significantly lower for flies from T2 (middle time-point) (median = 60.38 minutes, interquartile range (IQR) = 4.55, N = 72) in comparison to early (T1) (median = 73.22 minutes, IQR = 3.07, N = 72) (estimate = −13.151, t = −27.115, P < 0.0001) and late (T3) (median = 65.39 minutes, IQR = 3.71, N = 72) (estimate = 4.275, t = 8.814, P < 0.0001) time-points. Variation in heat knockdown response is partially explained by variation in experimental cages too (df = 6, F= 42.12, P< 0.0001) (Fig. 5).

**Figure 5:**
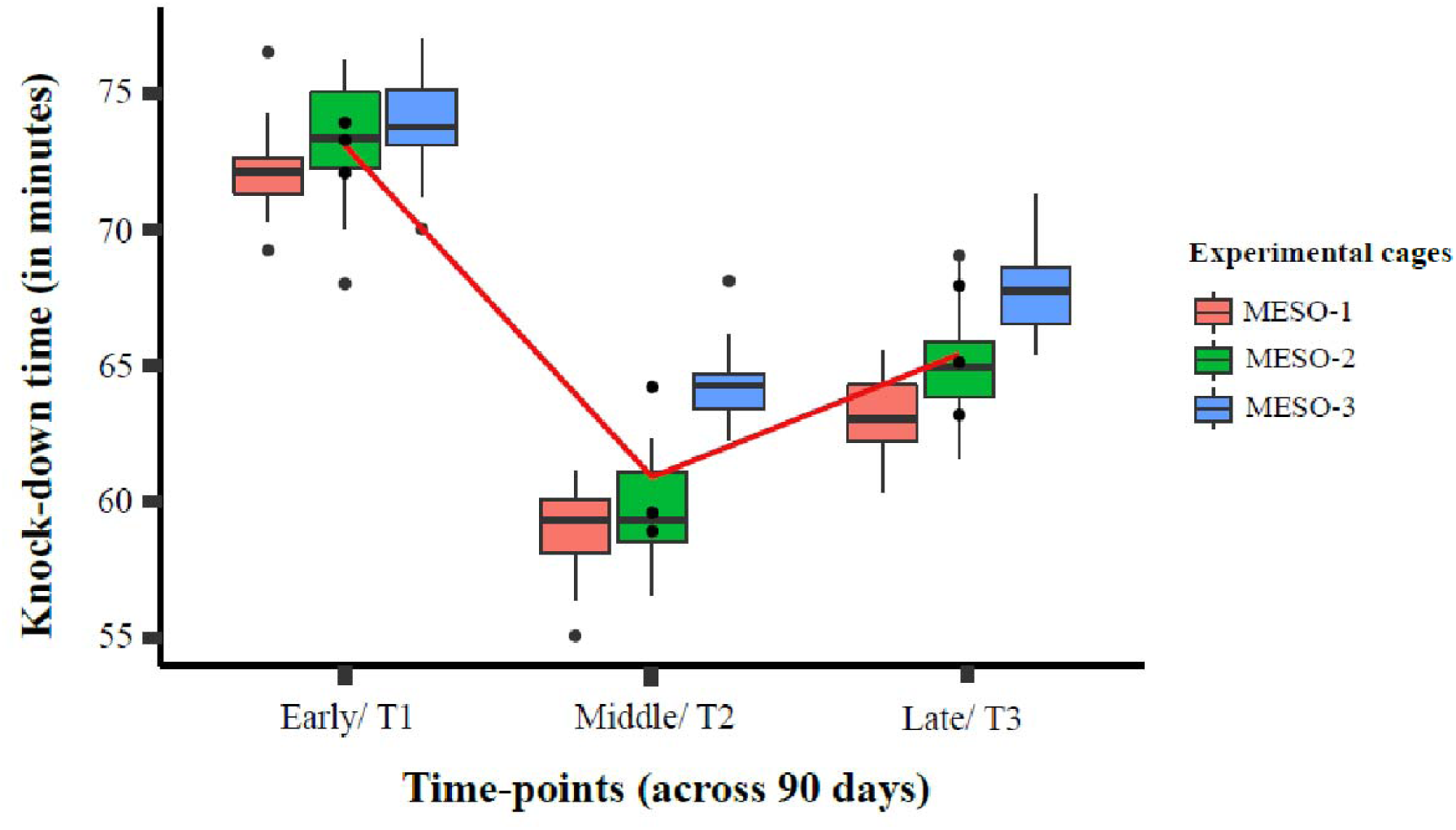
Box-plots representing variation in knock-down times in response to heat ramping assays, across three time-points (early, middle and late) from three replicate experimental cages. Flies from the middle time-point (45^th^ day) display the significantly lowest tolerance to increase in temperatures compared to the early (day 0) and late (90^th^ day) time-points.

## 4. Discussion

An organisms’ response to changing environmental conditions determines its fitness. Adaptations to rapid environmental fluctuations are known (Franks and Hoffmann 2012). Responses to rapid environmental change could be micro-evolutionary or mediated through phenotypic plasticity (Meyers and Bull 2002, Sgrò, Terblanche et al. 2016). Trait plasticity in response to climate change has been documented across diverse taxa, e.g. in birds (Charmantier, McCleery et al. 2008), mammals (Nussey, Wilson et al. 2007), reptiles (Leal and Gunderson 2012) and insects (Boggs 2016, Halsch, Shapiro et al. 2021). Still, studies investigating response to climate change have been attempted predominantly for temperate (Finn, Grattarola et al., Feeley, Stroud et al. 2017, Sheldon 2019) rather than tropical species (especially south-east Asia) (but see (Parkash, Tyagi et al. 2005, Rajpurohit, Zhao et al. 2017)). *D. melanogaster* originating from sub-Saharan Africa showing cosmopolitan distribution worldwide (David and Capy 1988, Lachaise, Cariou et al. 1988, Sprengelmeyer, Mansourian et al. 2019) has been investigated in recent years for responses to climate changes (Hoffmann, Hallas et al. 2003). The role of natural selection in latitudinal clinal variation in *D. melanogaster* across broad demographic zones of tropics and temperate regions is known (Kolaczkowski, Kern et al. 2011). However, genomic analyses for climatic adaptation have been studied in North American (Fabian, Kapun et al. 2012), European (Božičević, Hutter et al. 2016) African (Pool, Corbett-Detig et al. 2012, Sprengelmeyer, Mansourian et al. 2020) and Australian (Hangartner, Hoffmann et al. 2015) *D. melanogaster* populations. Parallel adaptations to cold climatic conditions are recorded for fly populations from Africa, Europe and North America (Božičević, Hutter et al. 2016, Pool, Braun et al. 2017). Here, we investigated the response of *D. melanogaster* collected from colder regions of the western Himalayas to a growing summer season in outdoor cages (mesocosm). Thus, our population was sourced from a high altitudinal zone (where maximum temperatures do not escalate above 15^0^C) hence cold adapted, but subjected to a warm/ hotter summer season (maximum temperatures can soar to 45^0^C). We expected thermos-sensitive genes to be upregulated in response to the growing season. However, we found no such pattern. Laboratory experiments demonstrate that *D. melanogaster* flies adapting to fluctuating thermal regimes increase their thermal breadth. Thus, *D. melanogaster* lines exposed to alternate cycles of high and low temperatures exhibited a generally tolerance to both cold and heat extremes (Tobler, Hermisson et al. 2015). Cold hardened flies survived better than non-hardened flies after a heat-stressor treatment. Tolerance to heat could be due to the activation of heat-shock proteins (Hsp70) which could have been inactive at low temperatures initially (Sejerkilde, Sørensen et al. 2003). Contrarily, cold adapted population from the Himalayas did not exhibit strong tolerance to summer conditions (Fig. 5) probably because selection for a colder climate could have reduced the physiological capacity to tolerate higher temperatures.

We used whole genome sequencing and filtered common SNPs for TRGs across the three time-points (T1 to T3). Results demonstrate considerable allelic variation across the three time-points. Temperature would increase from T1 to T2 and then decrease from T2 to T3. We therefore surmise that allelic variation could execute through physiological plasticity in response to temperature extremes in this tropical species. Higher allelic variation was observed in the first two time-points in comparison to the third (T3), where temperature could plateau due to the onset of monsoon. Further studies could determine if the observed allelic variation corroborates positively with survival and fitness in the peak hot season. In temperate orchards, considerable allelic variation with respect to seasons (winter versus fall) has been noted (Bergland, Behrman et al. 2014) maintained through balancing selection. However, in this case, considerable allelic variation for temperature responsive genes was seen within a span of 90 days (corresponding to peak summer season). It therefore remains to be analyzed if selection is indeed strong, relaxed or balancing through seasonal oscillations like temperate counterparts.

While the mechanistic basis of temperature regulating TRGs is still being explored (Singh, McGoldrick et al. 2019), our study which reveals novel SNP’s in *Drosophila* TRGs opens up new avenues for testing thermal acclimation hypotheses. The pleiotropic effect of temperature sensation on other functions can be seen through the association of different pathways integrated through the protein interaction map. Nonetheless, genes associated are novel and not typical of those associated with thermal acclimation, e.g. heat shock proteins (Sørensen and Loeschcke 2007).

We found two independent lines of evidence: First, a large number of kinases and phosphorylases are associated with stress, possibly linked to heat/temperature as evident from the protein interaction map (Fig. 4). Second, the large number of key regulators and transcription factors that are associated with cell proliferation and growth. Underlying these, *Myc* emerged as a key gene with the interactants possibly associated with genetic arrangements, and most importantly reducing the mutational load or burden. This is also in agreement with the fact that the *Myc* plays a key role in insufficiency of haplotypes reducing the mutation load which further is the key factor for extended lifespan or pro-aging (Morrison, Murakami et al. 2000, Greer, Lee et al. 2013). On the other hand, phosphatidylinositol 5-phosphate 4-kinase (PIP4K) is known to regulate the growth during fly development (Gupta, Toscano et al. 2013). While its activity is implicated in cellular responses, regulating the growth factors, the protein interaction networks clearly reveal that there are transient mutations underpinning the changes in heat/temperature stress as these changes could be because of the activity of downstream kinases. We envisage that the pathways associated with PIP4K switch signals modulating the strength of a signal. Taken together, the kinases and phosphatases contribute to the outcome or ability to acclimate the flies in these conditions. As they adapt, essential regulatory processes and changes mediating phosphorylation and kinases are known. Further, our genomic sequencing has emerged as an approach to understand such “adaptive loci’’ influencing these signaling pathways.

Mesocosm experiments as ours closely resemble natural conditions are hitherto reported only in temperate populations (e.g. (Rudman, Greenblum et al. 2022)) but not for tropical species. Ours is therefore the first study to track genomic genetic variation at more realistic and natural time-scales in a tropical high altitudinal fruit fly species. We considered a time frame where summer temperature steadily increases, peaks and then wanes. Our results from heat ramp assay corroborate this, in that increase in temperatures of the external environment (cages) also altered the thermal sensitivity. Thus, the peak in the temperature during T2 (Fig. 1) also corresponded with flies from T2 being the most sensitive to thermal stress and hence the faster knock-down time in T2 compared to T1 and T3 (Fig. 5). However, further studies could ascertain if seasonal variation in alleles related to TRGs are connected/ pleiotropic with fitness-related traits. Thus, in the temperate regions, diapause associated genes are known to be up and down regulated corresponding to winter and fall seasonal variation, respectively (Zhao, Bergland et al. 2016) with outcomes in fecundity (ovariole size (Schmidt, Matzkin et al. 2005)). In tropical scenarios, where adaptations like diapause are less known, one could expect sensitivity towards thermal extremes to evolve through alternate gene regulatory networks. Since *Drosophila melanogaster* originated from Africa (David and Capy 1988) and then dispersed to other continents, reaction norms for temperature tolerance could be expected to be broadly similar across the tropics than across temperate regions. Surprisingly, our study populations from higher altitudes (Rohru, western Himalayas) exhibited limited tolerance to higher temperatures, thus re-affirming the speculation that tropical species are indeed sensitive to higher temperature thresholds. It remains to be tested if differential response to temperature extremes does exist across altitudinal populations. Given clinal variation for multiple traits noted for Drosophilids from the Indian subcontinent (Rajpurohit, Zhao et al. 2017) (Rajpurohit, Oliveira et al. 2013, Rajpurohit, Zhao et al. 2017), exploring thermal tolerance and associated genomic variation is an interesting research avenue. Further investigations could involve understanding chromosomal inversion clines which have been linked with local adaptation in *D. melanogaster* (Kapun, Fabian et al. 2016, Kapun and Flatt 2019). Inversion clines are common across all continents where *D. melanogaster* has dispersed, yet they have been less addressed from south-east Asia (Das and Singh 1990) where the inversion frequency is speculated to be low (Glinka, Stephan et al. 2005). Surprisingly, inversion polymorphisms linked with climate adaptation were found to be less frequent at higher altitudes than lowlands in Africa populations (Aulard, David et al. 2002, Pool, Braun et al. 2017). Does the ancestral (sub-Saharan) derived inversion polymorphism confer selective advantage to tropical high altitudinal *D. melanogaster*? Whether south-east Asian flies evolved alternative loci/ gene networks as our study reveals needs further exploration.

## Conclusions

Tropical species are likely to be affected by global increase in temperature since the thermal range to which they could respond is narrow compared to their temperate counterparts. Our study hints that the narrow temperature tolerance of tropical species could be linked with alternate molecular pathways for thermal sensitivity. We found a considerable number of newer candidate alleles whose frequencies are significantly shifting as temperature increases over the growing season. Our study highlights that genomic signatures could be instrumental to understand organismal responses to changing environmental conditions. Our findings of allelic variation have implications not only in tracking fitness outcomes across seasons but also how organisms are evolved in the face of rapid climate change. Future work could combine molecular and field based approaches for a comprehensive understanding of climate mediated changes.

